# escheR: Unified multi-dimensional visualizations with Gestalt principles

**DOI:** 10.1101/2023.03.18.533302

**Authors:** Boyi Guo, Louise A. Huuki-Myers, Melissa Grant-Peters, Leonardo Collado-Torres, Stephanie C. Hicks

## Abstract

The creation of effective visualizations is a fundamental component of data analysis. In biomedical research, new challenges are emerging to visualize multi-dimensional data in a 2D space, but current data visualization tools have limited capabilities. To address this problem, we leverage Gestalt principles to improve the design and interpretability of multi-dimensional data in 2D data visualizations, layering aesthetics to display multiple variables. The proposed visualization can be applied to spatially-resolved transcriptomics data, but also broadly to data visualized in 2D space, such as embedding visualizations. We provide an open source R package `escheR`, which is built off of the state-of-the-art `ggplot2` visualization framework and can be seamlessly integrated into genomics toolboxes and workflows.

**Availability and implementation:** The open source R package `escheR` is freely available on Bioconductor (bioconductor.org/packages/escheR).

## 1 Introduction

Visualization is an indispensable component of data analysis, providing clarity that connects quantitative evidence to key conclusions [6]. In biomedical research, visualization receives growing recognition as essential: many scientists rely on visualization to complete their cognitive process from analysis to insight, including analytic validation of automated pipelines and scientific communication [18]. However, an important challenge in biomedical research is how to visualize increasingly complex, multi-dimensional data [19].

Here, we focus on two types of visualizations in biomedical research, but note that the proposed ideas could be extended beyond these applications: (i) embedding visualizations, which project data into some low-dimensional embedding or mathematical space (e.g. Principal Components Analysis [11], *t*-distributed Stochastic Neighbor Embedding (*t*-SNE) [28], or Uniform Manifold Approximation and Projection (UMAP) [3]) and (ii) *in situ* visualizations [5, 16, 18], which aim to visualize molecules captured from *in situ* imaging or sequencing technologies where *in situ* refers to ‘in its original place’. Both of these visualizations represent data in a 2D space and are motivated by recent advances in experimental technologies that profile molecules, including DNA, RNA, and proteins, at a single-cell or spatial resolution [13, 17]. Some most popular technologies include single-cell/nucleus RNA-sequencing (sc/snRNA-seq) [1] and *in situ* spatially-resolved transcriptomics [15].

A common and fundamental challenge with both of these visualizations is how to visualize multi-dimensional information in a 2D space. For example, in *in situ* visualizations, we often want to create a spatial map to visualize a continuous (e.g. gene expression) or discrete (e.g. cell type or spatial domain) variable representing molecular information in the original spatial location. However, it is challenging to simultaneously visualize multi-dimensional data, such as information from disparate data domains (such as expression domain and spatial domain) or disparate data modalities (such as transcriptomics and proteomics) in the same plot. Currently, best practices for this include making two different plots displayed side-by-side (**Figure 1A-B**), one for gene expression and one for spatial domains. This creates cognitive gaps on how to associate the disparate information or how to interpret the biological findings of this multi-dimensional information regarding their (micro-)environment or colocalization. While interactive visualizations [14, 21, 26] have the potential to mitigate this challenge, they are infeasible for scientific communications in static media, such as printed work. Developing a static and unified visualization that enables the simultaneous display of multiple dimensions of information is crucial for biomedical research.

**Figure 1:**
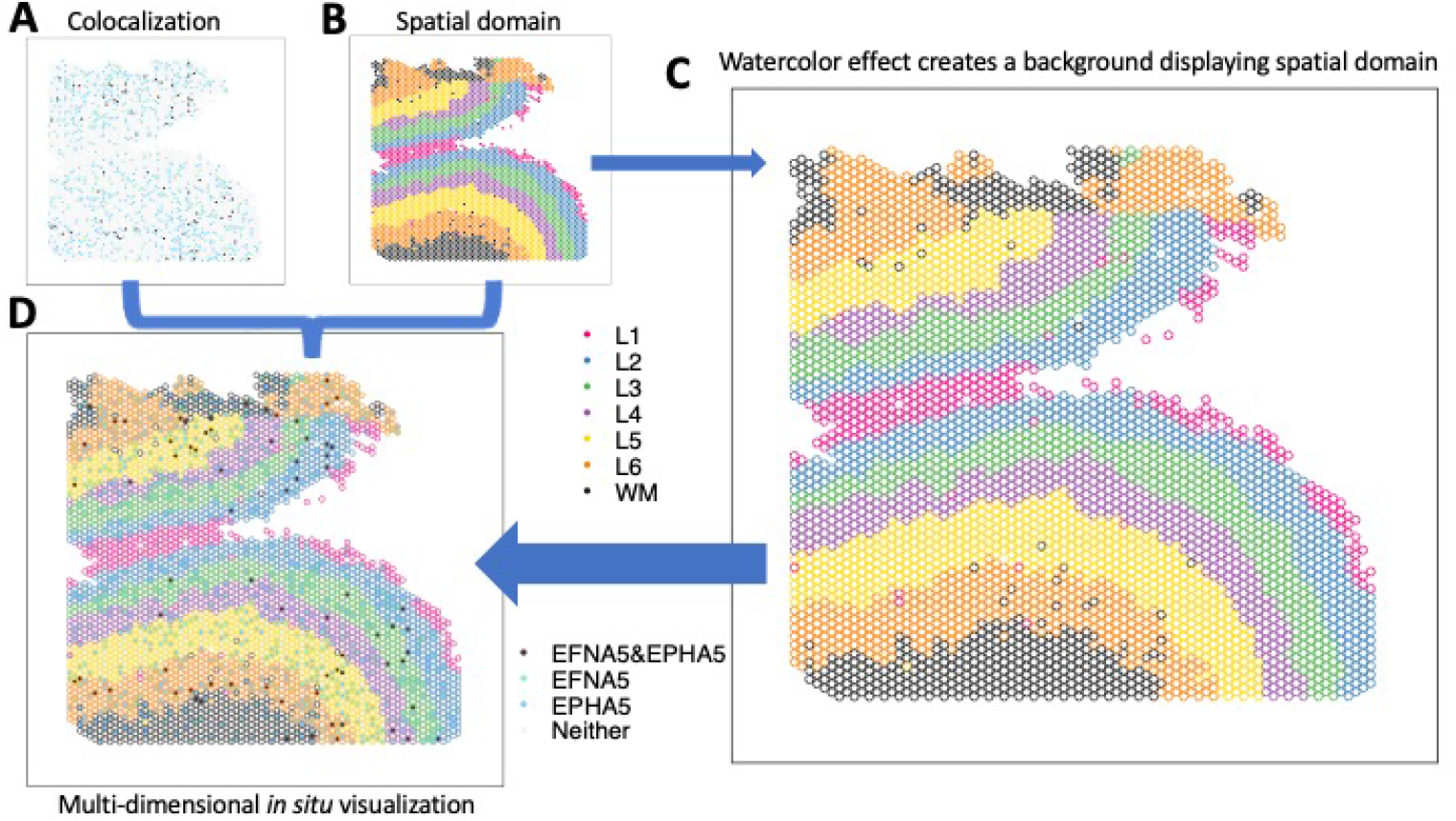
escheR enables multi-dimensional spatial visualizations following the Gestalt principles of design. (**A-B**) The traditional visualization displays the colocalization plot of the expression of two genes *EFNA5* and EPHA5 (**A**) and the spatial domains from the dorsolateral prefrontal cortex in postmorteum human brain [12] (**B**) side-by-side, creating challenges to cognitively connecting colocalization status to spatial domains. (**C**) The watercolor effect enables displaying spatial domains by color-coding only outlines of circles. (**D**) escheR enables the multi-dimensional *in situ* visualization that simultaneously displays the cortex layers and the colocalization status, substantially improving interpretability.

## 2 Results

To address these challenges, here we leverage the Gestalt (German for “unified whole”) principles for design [20, 27] as a way to visualize multi-dimensional data in 2D visualizations. We focus on the two types of data visualizations previously introduced that are widely used in biomedical research: (i) embedding visualizations and (ii) *in situ* visualizations. We provide an R package, `escheR`, implementing these ideas, which is built on the state-of-the-art data visualization framework `ggplot2` in the R programming language. Finally, we comment on how these ideas could be extended to other types of visualization in biomedical research.

### 2.1 Multi-dimensional 2D visualizations with ggplot2 and Gestalt principles

Gestalt principles [20, 27] refer to a set of rules describing how humans perceive and interpret visual information and are commonly applied in art and designs. Developed in the 1920s by German psychologists Max Wertheimer, Kurt Koffka and Wolfgang Kohler, these principles help humans perceive a set of individual elements as a whole.

Here, we leverage the principles to be able to visualize multi-dimensional data in a unified 2D plot. Our approach is to use the state-of-art data visualization framework `ggplot2` [29] following the Grammar of Graphics [30] and map individual variables to different aesthetics to simultaneously display disparate variables. Specifically, we apply the figure-ground segmentation [22] in displaying two variables: one variable (e.g. expression) can be plotted as color-filled circles, serving as the *figure*; one variable (e.g. spatial domains) can be plotted as the backgrounds of the circles, creating a *ground* for the figure. In practice, we use the combination of color and fill=“transparent” to create the background layer and fill to create the figure layer. When necessary to display an additional layer for a third variable, shape can be used to add symbols such as cross (+) and asterisk (*) to highlight in the spatial map.

For adjacent circles with limited space between them to display the background color, we use an economic implementation, colored outlines for these circles (**Figure 1C**), inspired by watercolor effect [23, 2 4]. Watercolor e ffect describes the phenomenon in visual perception that surface color arises from thin boundaries and hence is applied here to perceive the background color in tight space. Overall, the figure-ground segmentation creates two isolated layers in visual perception to display the two variables while maintaining the relative spatial relationship serving as a reference between the two. In addition, other fundamental principles [27], such as proximity, similarity, continuity, and closure, incentivize the brain to group elements and dimensions in the visualization, guaranteeing an integrative perception of the complex multi-dimensional spatial map.

Here, we provide an open-source package called `escheR` (named after the graphic artist M.C. Escher) in the R programming language [25], leading to a simplified interface to navigate the implementation of the multi-dimensional visualization in 2D space. By adapting `ggplot2` standard, `escheR` can be seamlessly integrated into many popular spatial resolved toolboxes, such as `SpatialLIBD` [21], `Seurat` [9], `Giotto` [5] to name a few, and allow further theme customization with ease.

Next, we give two use cases to exemplify some utility of the proposed spatial visualization: (i) the spatially differential gene colocalization in the human dorsolateral prefrontal cortex using spatial transcriptomics data [12]; (ii) multi-dimensional UMAP highlighting differential gene expression in data-driven cell clusters [7].

### 2.2 Multi-dimensional *in situ* visualization

In a recent study investigating the molecular organization of human dorsolateral prefrontal cortex [12], two schizophrenia risk genes, membrane-bound ligand ephrin A5 (*EFNA5*) and ephrin type-A receptor 5 (*EPHA5*), were identified to colocalize via cell-cell communication analysis. In addition, data suggested Layer 6 was the most highly co-localized layer compared to other cortex layers. To visually examine the inference, we applied `escheR` to create a multi-dimensional *in situ* spatial map that simultaneously exhibits the cortex layers (displayed with color-coded spot outlines) and the categorized colocalization status of genes *EFNA5* and *EPHA5* (displayed with color-coded spot fill). Compared to the traditional visualization where the cortex layers and the colocalization status are visualized in two side-by-side figures (**Figure 1A-B**), our proposed visualization (**Figure 1D**) enables directly mapping colocalization status to the spatial domain, simplifying the perception of two sources of information and allowing cognitive comparison across cortex layers.

### 2.3 Multi-dimensional embedding visualizations

The application of the proposed framework is not limited to *in situ* visualizations of spatially-resolved transcriptomics data. It is broadly applicable to data mapped to any 2-dimensional coordinate system to simultaneously display multiple variables. Such systems include euclidean space (including spatial coordinate as a special case) and data-driven embedding space, for example, UMAP and *t*-SNE. To demonstrate, we applied the proposed visualization to address the challenge of simultaneously displaying cluster membership and gene expression in a single-cell UMAP plot. To address the overplotting problem, previous work proposed to apply hexagonal binning strategy to display the gene expression [7]. Here, the color-coded convex hulls are used to annotate different clusters of cells (**Figure S1A**). However, the convex hulls create substantial overlapping areas, creating confusion when interpreting cluster memberships of hexagons in the overlapping areas. To improve the interpretability of the visualization, we replace the convex hulls with color-coded hexagons boundaries (**Figure S1B**) to avoid possible membership confusion. We note that our contribution to improving the visualization is easily implemented without any modification of `schex` as both are built upon the Grammar of Graphics [30] standard.

## 3 Discussion

Here, we propose an innovative multi-dimensional spatial visualization that simultaneously displays multiple variables in a 2D coordinate system. Specifically, our design leverages Gestalt principles from visual perception to create multiple visual dimensions in a spatial map by iterative layering aesthetics. Built upon `ggplot2`, we provide an open-source R package `escheR` that is seamlessly compatible with popular spatially-resolved transcriptomics and single-cell data analysis toolboxes.

Adding a third dimension to 2D plots has been a long-standing challenge in visualization [19]. Our proposal addresses this fundamental challenge by introducing simple but effective design principles. These principles lead to visually easier-to-interpret graphics compared to 2D-color gradients and geometry annotations (**Figure S1**). Unlike computer-based interactive visualizations, the proposed visualization is free from any platform and technology restriction, creating an accessible and economical solution. In addition, the proposed visualization is easily scalable and hence can be applied to all types of spatially resolved data. Combining with a binning strategy, similar to `schex` [7], to avoid possible overplotting, the proposed visualization can be also applied to visualized image-based spatially resolved data [4], in addition to aforementioned spot-based spatially resolved data (**Figure 1**). `escheR` could also be applied with kernel gene-weighted density plots from `Nebulosa` [2] and other ggplot2-based visualizations.

Beyond the scope of biomedical research, the proposed visualization can be broadly translated to any visual analytic highlighting differentiation with respect to another measurement(s). To name a few, such visual analytics include examining differential tests, explaining clustering, and visualizing subgroups. However, one of the most rewarding fields to apply the proposed visualization is the rapidly expanding field of biomedical multi-omics research [10], where connecting different omics (data modalities) is the fundamental goal and hence greatly appreciating innovative multi-dimensional visualization.

In summary, we propose a novel multi-dimensional visualization, implemented in an R package `escheR`, to address the simultaneous exhibition of multiple variables in 2D plots. The proposed visualization can be broadly applicable to the visual analytics of growingly complex biomedical data and beyond.

## Abbreviations

PCA: principal component analysis
*t*-SNE: *t*-distributed stochastic neighbor embedding
UMAP: uniform manifold approximation and projection
scRNA-seq: single-cell RNA-sequencing
snRNA-seq: single-nucleus RNA-sequencing
SRT: spatially-resolved transcriptomics
EFNA5: membrane-bound ligand ephrin A5
EPHA5: ephrin type-A receptor 5
DLPFC: dorsolateral prefrontal cortex

## Author contributions

- Boyi Guo: Conceptualization, Methodology, Software, Validation, Formal analysis, Investigation, Data Curation, Writing, Visualization
- Louise A. Huuki-Myers: Conceptualization, Software
- Melissa Grant-Peters: Conceptualization
- Leonardo Collado-Torres: Conceptualization, Software
- Stephanie C. Hicks: Conceptualization, Resources, Writing -Review & Editing, Visualization, Supervision, Project administration, Funding acquisition

## Declarations

### Ethics approval and consent to participate

Not applicable.

### Availability of data and materials

The spatial transcriptomics dataset was obtained from `spatialLIBD` (research.libd.org/spatialLIBD). The UMAP example follows the ‘using_schex’ vignette in the `schex` package (bioconductor.org/packages/schex). The code that generates these figures is deposited at github.com/boyiguo1/Manuscript_escheR (Zenodo DOI: 10.5281/zenodo.7915970). The open source software package `escheR` available in the R programming language is freely available on GitHub (github.com/boyiguo1/escheR) and Bioconductor (bioconductor.org/packages/escheR).

## Acknowledgments

We would like to acknowledge Nicholas J. Eagles, Kristen R. Maynard, Mina Ryten, Leon Di Stefano, and Lukas M. Weber (appearing in alphabetic order of last name) for their helpful comments, feedback and suggestions on `escheR` functionality. Nicholas J. Eagles and Kristen R. Maynard are employed by the Lieber Institute for Brain Development; Mina Ryten is employed by University College London; Leon Di Stefano and Lukas M. Weber are from the Johns Hopkins Bloomberg School of Public Health, Department of Bio-statistics.

## Supplementary Materials

### Supplemental Figures

**Supplementary Figure S1:**
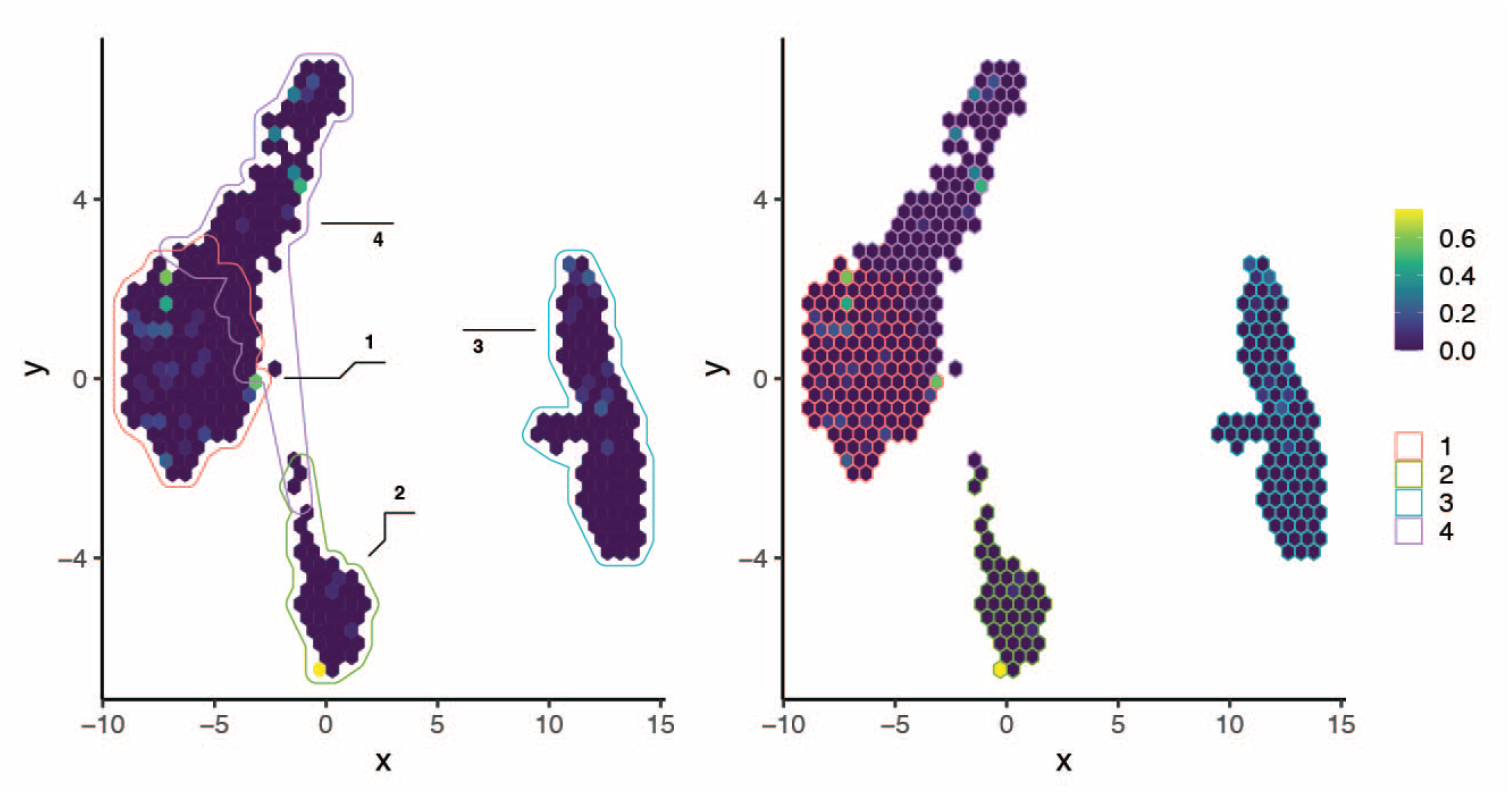
escheR enables multi-dimensional embedding visualizations. The gene expression of *POMGNT1* among peripheral blood mononuclear cells [8] under the UMAP representation. (**A**) The schex R/Bioconductor package uses color-coded convex hulls to annotate data-driven cell types, creating confusion when interpreting hexagons in overlapping hulls. (**B**) escheR plots hexagon-specific membership to improve interpretability.

