## Supplementary Materials for "escheR: Unified multi-dimensional visualizations with Gestalt principles"

##### Contents

1. **Supplemental Figures S1.**

### Supplemental Figures

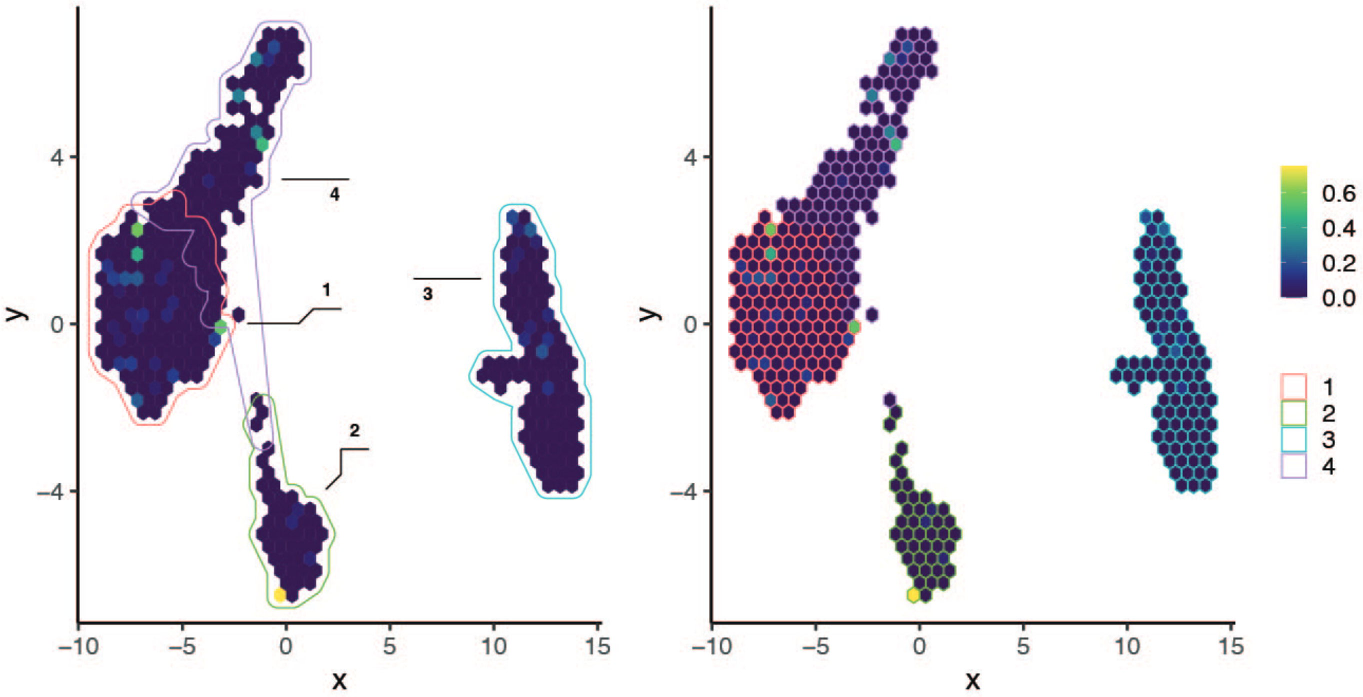


Supplementary Figure S1: **escheR enables multi dimensional embedding visualizations**. The gene expression of *POMGNT* among peripheral blood mononuclear cells [29] under the UMAP representation. (**A**) The `schex` R/Bioconductor package uses color-coded convex hulls to annotate data-driven cell types, creating confusion when interpreting hexagons in overlapping hulls. (**B**) `escheR` plots hexagon-specific membership to improve interpretability.
